# Transcranial direct current stimulation does not improve performance in a whole-body movement task

**DOI:** 10.1101/2021.01.25.428100

**Authors:** Harriet Caesley, Isabella Sewell, Nikita Gogineni, Amir-Homayoun Javadi

**Author notes:** **Corresponding author:** Amir-Homayoun Javadi, Address: School of Psychology, Keynes College, University of Kent, CT2 7NP, Canterbury, Kent, UK.

## Abstract

Research has investigated the use of non-invasive brain interventions, such as transcranial direct current stimulation (tDCS), to enhance motor learning and rehabilitation. Much research has shown that tDCS improves motor learning and that bilateral tDCS is more beneficial than unilateral tDCS in improving motor learning. However, past research has primarily utilised simple motor tasks in measuring motor skill learning. These are not ecologically reliable as whole-body movement is required for everyday activities. This study involved two experiments. Each experiment involved participants learning 12 Ballroom and Latin dance moves whilst undergoing tDCS. All participants underwent three sessions of tDCS, (unilateral, bilateral and sham), over three consecutive days. Participants in the first experiment (n=30) had stimulation to the primary motor cortex (PMC) and those in the second experiment (n=31) had stimulation to the dorsolateral prefrontal cortex (DLPFC). In each experiment, a baseline was taken before the training sessions and two outcome measures were taken; a day after the last training session and two weeks later. In each testing session participants’ dance ability was measured. Our results showed that bilateral tDCS impaired performance in both experiments. Unilateral stimulation impaired performance in the first experiment, and did not significantly improve performance any better than the sham stimulation in the second experiment. These results suggest that task complexity plays a crucial role when tDCS procedures are used to modulate motor performance and highlights possible limitations of tDCS in practice.

## 1 Introduction

Transcranial electrical brain stimulation, a noninvasive and painless brain stimulation method, has been shown to be effective in cognitive enhancement (Parking et al, 2015). It has also been shown to be beneficial in improving simple motor movements, such as; serial reaction time tasks (Giustiniani et al, 2019), upper limb rehabilitation in stroke patients (Arnao et al 2019) and finger tapping in individuals with neurological disabilities (Bolognini et al. 2011). However, research looking at the effect of transcranial electrical brain stimulation on whole-body movement is minimal. Therefore, this study aims to look at different protocols of tES and its ability to impact upon whole-body movement.

One popular technique of transcranial electrical brain stimulation is transcranial direct current stimulation (tDCS). Its popularity being the result of its affordability and easy to use nature (Reis & Fritsch, 2011). tDCS involves the transmission of low intensity currents to specific areas of the brain via electrodes attached to the scalp. Generally, it is assumed that an anodal electrode will increases excitability whereas a cathodal one will decrease excitability (Bindman, Lippold & Redfearn, 1964). tDCS effects can also last after stimulation ceases, and can potentially last up to 90 minutes (Nitsche & Paulus, 2001). Research on the effects of tDCS on minor movements is plentiful (Apšvalka, Ramsey & Cross, 2018; Hashemirad, Zoghi, Fitzgerald & Jaberzadeh, 2016; Moura et al. 2017; Rocha et al. 2016; Srirman et al. 2014). Minor movements occur in the wrist, hands, fingers, feet and toes, and typically only involve movements with one goal. Positive effects of tDCS have been found in studies looking at; motor adaptation (Weightman, Brittain, Punt, Niall & Jenkinson, 2020), throwing tasks (Jackson et al 2019), serial reaction time finger tapping tasks (Ehsani, Bakhtiary, Jaberzadeh, Talimkhani & Hajihsani 2016; Talimkhani et al 2019) and balance tasks (Kaminski et al 2016; Zandvliet, Meskers, Kwakkel & van Wegen 2018). Additionally, similar positive effects have been found in patients suffering from neurological injuries and illnesses, such as in stroke (Rocha et al, 2016; Allman et al, 2016) and Parkinson’s disease (Kami, Sadler, Nantel and Carlsen, 2018).

All studies above used a unilateral stimulation configuration. In these studies, only one hemisphere was actively stimulated. Stimulation of both hemispheres is known as bilateral stimulation and research suggests this also positively impacts motor learning (Bologni et al. 2011; Waters-Metenier, Husain, Wiestler & Diedrichsen, 2014). Furthermore, coupled with physical therapy bilateral stimulation has proven to be beneficial in treating individuals with motor disabilities such as those with stroke (Lindenberg, Renga, Zhu, Nair & Schlaug, 2010; Bologni et al., 2011). Moreover, it has been suggested that the effects of bilateral stimulation are stronger and last longer than unilateral stimulation (Sehm, Kipping, Schäfer, Villringer and Ragert, 2013; Vines, Cerruit and Schalug, 2008). The long-lasting effects of bilateral tDCS make this a more appropriate technique in the treatment for individuals with motor disabilities. A possible explanation of this superior effect, Waters, Wiestler and Diedrichsen (2017) have suggested that higher efficacy for bilateral stimulation is due to electrical currents running transversely, subsequently increasing the plasticity in both primary motor cortices (Waters, Wiestler and Diedrichsen, 2017; Lindenberg, Sieg, Meinzer, Nachtigall & Flöel, 2016).

While past research has suggested beneficial effects of tDCS in paradigms for simple-body movements, the application of tDCS and whole-body motor movement learning has been neglected greatly (Kaminski et al. 2013; Kaminski et al. 2016; Steiner et al. 2016). By identifying the relationship between the two, results can be used to provide a more conclusive evaluation of the effects of tDCS on motor learning. Additionally, the results could also be implemented into treatment programmes to help the motor recovery of individuals who have suffered from illnesses and diseases. Therefore, in this study we investigated the effect of different protocols of tDCS on simple dance moves, which is a form of whole-body movement.

## 2 Experiment 1 – primary motor cortex stimulation

Previous research has shown that the primary motor cortex (PMC) is active during a motor task (Dushanova & Donoghue 2010; Honda et al 1998; Kakei, Hoffman & Strick 1999; Muellbacher et al, 2000). The PMC is important in movement and more specifically motor learning because it facilitates; motor adaptation (Ehsani et al. 2016), skilled voluntary movements (Kida & Mitsushima, 2018) and fast online performance improvement (Karok, Fletcher & Whitney 2017). Consequently, tDCS of the PMC also enhances movement (McCambridge, Bradnam, Stinear & Byblow 2011), motor learning (Ciechanski & Kirton 2016; Karok et al 2017; Yamaguchi et al 2020), motor sequence learning (Hashemirad et al 2016; Stagg et al 2011) and motor learning of novel skills (Dumel et al, 2018).

We hypothesised that active anodal-tDCS over the right-PMC will be more beneficial than sham stimulation at improving an individual’s ability to learn dance moves. Furthermore, we hypothesised that bilateral anodal tDCS over both motor cortices will be more advantageous than unilateral stimulation over the right-PMC at improving an individual’s ability to learn dance moves.

### 2.1 Method

#### 2.1.1 Participants

The sample included 30 participants (29 females, age mean[SD] = 18.97[1.26] years old). None of the participants had experience with Latin or Ballroom dancing, as indicated in a pre-study questionnaire. All participants gave written informed consent. The study was approved by the local ethics committee in the School of Psychology at the University of Kent.

#### 2.1.2 Materials

*Dance videos and scoring* – Twelve dance videos were recorded. For the details of the dance moves please see the supplementary document. Each move was performed by a male and a female dancer, so to match the participants’ gender. Criteria for the scoring stimuli was created by an experienced Latin and Ballroom dancer. Participants were evaluated on posture, size of movement, timing, arms, legs and overall performance ability. Criteria for arms, legs and overall performance was different depending on the move performed. Please see the supplementary for the details. Scoring was done by the two experimenters (one experienced Ballroom and Latin dancer and one with limited experience), there was a significant positive relationship between both experimenters (r(46) = .865, p < 0.001), indicating that there is a strong inter-rater reliability between the marking scores of each experimenter.

*Transcranial Direct Current Stimulation (tDCS)* – One or two tDCS (NeuroConn, Germany) stimulators were used with current amplitude of 1.5mA and 10 seconds ramp up and down. 1.5mA was used as previous research has suggested that 2mA does not improve learning (Hoy et al. 2013; Parkin, Bhandri, Glen & Walsh, 2019) and 1.5mA was more effective in a motor learning task than 1mA (Cuypers et al. 2013). Stimulation was applied to either the right primary motor cortex (Unilateral stimulation) or both primary motor cortices (Bilateral stimulation). According to the international EEG 10-20 system, the active anodal electrode was placed on C4 (Unilateral stimulation), and C3 and C4 (Bilateral stimulation) (Jasper, 1958). Depending on the stimulation protocol one or two cathode electrodes were placed on the upper arms, contralateral to the side of corresponding anode electrode, Figure 1. Electrodes were 5×5cm^2^, and were soaked in salt-water solution. For unilateral and bilateral stimulation protocols, stimulation was applied for five minutes before motor learning commenced, and then continued throughout the training paradigm for the maximum duration of 15 minutes. For sham stimulation, the placement of the electrodes was the same as unilateral stimulation, however stimulation was only applied for 20 seconds (10s ramp up and down).

**Figure 1.**
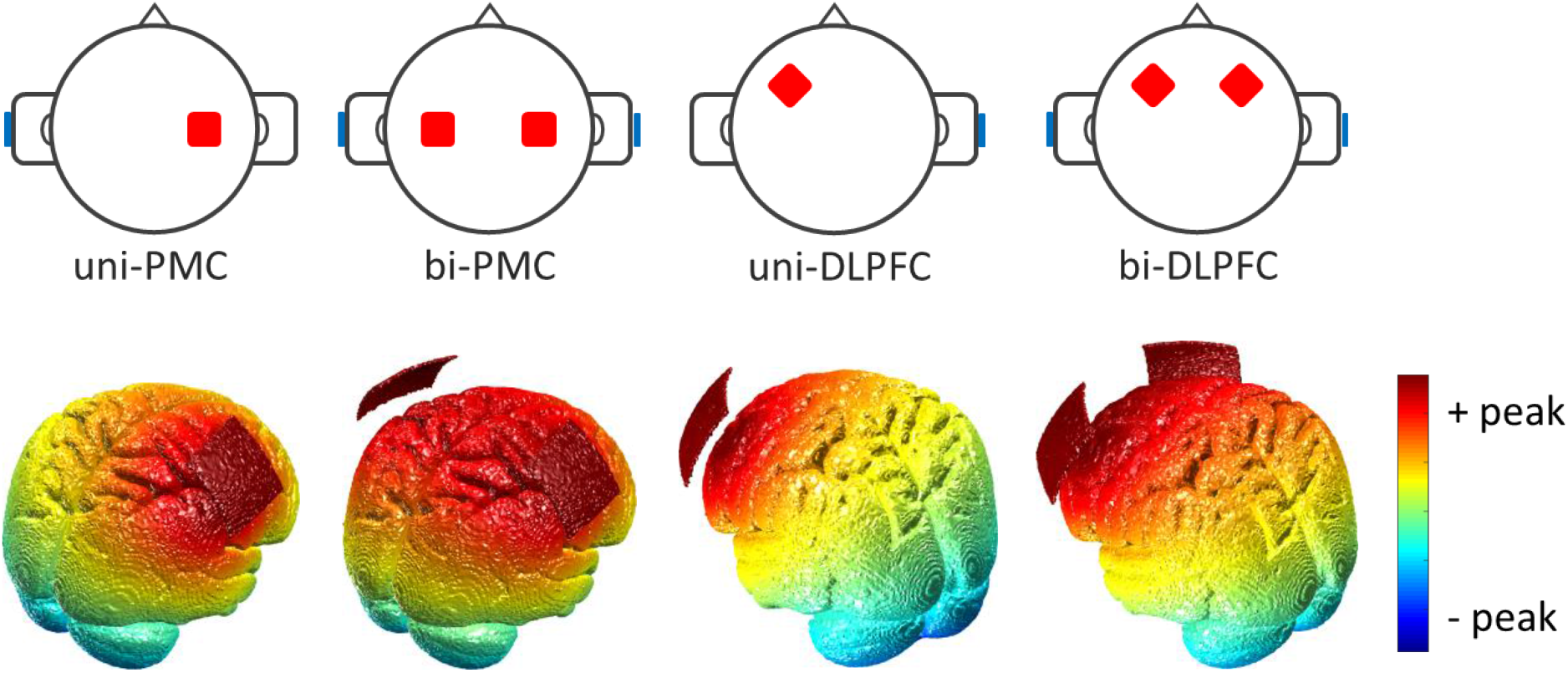
Placement of the electrodes (top panel) and simulation of the brain stimulation (bottom panel) (Huang et al, 2019) for Experiment 1 (stimulation of the primary motor cortex; PMC) and Experiment 2 (stimulation of the dorsolateral prefrontal cortex; DLPFC) using unilateral (uni-) and bilateral (bi-) electrode montage. Anode electrodes were placed on the target area and cathode electrodes were placed on the contralateral shoulder. Modelling was done using open-source software ROAST (Huang, Datta, Bikson, & Parra, 2019).

### 2.2 Design

A within-subjects design was used; participants were involved in all three conditions of the experiment. The independent variable had three levels (Bilateral/Unilateral/Sham). The dependent variable was percentage change in participants’ dancing ability from the baseline, scored based on the criteria. This was measured over three different days; baseline test, outcome measure one and outcome measure two.

### 2.3 Procedure

The study involved six sessions over six separate days; a baseline measure, three consecutive tDCS training sessions, an outcome measure one and an outcome measure two, see Figure 2.

**Figure 2.**
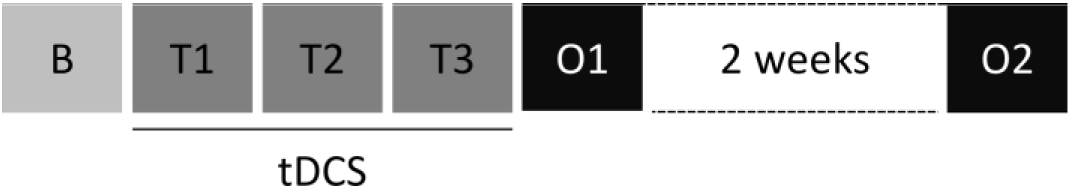
Summary of the procedure of the study. Participants took part in one baseline measure (B), followed by three training sessions (T1, T2 & T3) in which they received dance training in combination with different protocols of brain stimulation. Following the intervention, two outcome measures were conducted to investigate effects of training immediately following training (Post-intervention, O1) and lasting effects of training (Follow-up, O2).

The baseline measure involved the participant undertaking the dance test, where they were tested on all 12 moves. There were four ballroom moves, including Walz Box Step Forward, and Eight Latin moves, including Cha Cha New York and Samba Basic (See supplementary material for a full list). Participants would first watch the dance move performed twice and then they would perform the move to the best of their ability (without watching the video), this is when they would be marked on their dance ability. Once they have performed the move the test would proceed to the next move.

In each training session, the participant underwent a different type of brain stimulation; the stimulation type was randomised and the participant was blind to the type of stimulation being received. Whilst stimulation was running the participant practiced four of the original 12 moves. T1, T2 and T3 all involved a different set of 4 dance moves. The participant observed the move twice, danced along with the video four times, danced alone three times, danced along with the video twice more and then danced alone for a final time. After this sequence is completed the next move will be shown. In each training session, the dance moves were different and the order in which they were presented was random.

The outcome measure one and outcome measure two involved the same procedure as the baseline measure.

### 2.4 Analysis

To account for inter-subject variability, participants’ percentage change on sessions O1 and O2 was calculated based on their performance in the baseline session. A 3×2 repeated measure analysis of variance (rANOVA) was run with stimulation condition (Bilateral/Unilateral/ Sham) and session (Post-intervention/Follow-up) as within subject factors and performance percentage change as dependent variable.

### 2.5 Results

The rANOVA showed a significant main effect of stimulation condition (*F*(2,58)= 5.417, *p* = 0.007, *η_p_^2^* = 0.157) and session (*F*(1,29)= 37.491,*p* < 0.001, *η_p_^2^* = 0.564), but a non-significant interaction effect (*F*(2,58)= 0.230, *p* = 0.795, *η_p_^2^* =0.008). Post-hoc paired-sample t-tests showed significant difference between Sham and Unilateral (Sham mean[SD] = 42.73[0.18]%, Unilateral 37.52[0.15]%, t(29) = 2.213, p = 0.038, d = 0.404), and Sham and Bilateral (Bilateral 35.14[0.14]%, t(29) = 3.033, p = 0.006, d = 0.554), but no significant difference between Unilateral and Bilateral (t(29) = 0.921, p = 0.257, d = 0.168). See Figure 3 for a summary of the performance of the participants.

**Figure 3.**
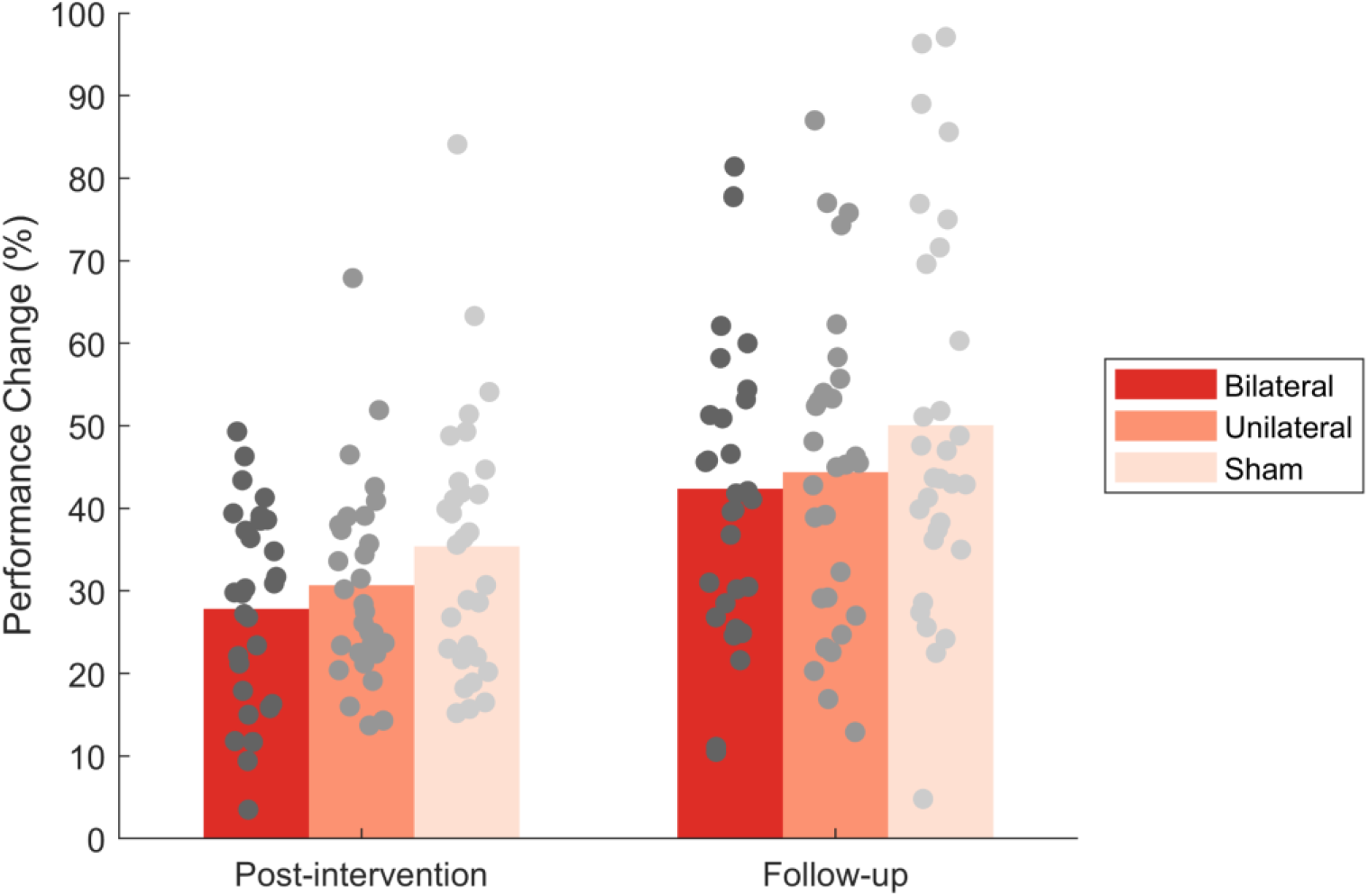
Performance percentage change from the baseline for the participants in different conditions and sessions in Experiment 1 with stimulation of the primary motor cortex. Sham stimulation showed significantly higher performance as compared to Unilateral (*p* = 0.038) and Bilateral *(p* = 0.006) stimulation conditions. No significant difference was found between Unilateral and Bilateral stimulation conditions *(p* = 0.257).

### 2.6 Summary

We investigated the effect of unilateral anodal stimulation over the right-PMC and bilateral anodal stimulation over both motor cortices on the learning of Ballroom and Latin dance moves. It was expected that active stimulation would be more effective than sham stimulation at improving dance performance and that specifically bilateral stimulation would be more beneficial than unilateral stimulation. However, contrary to our hypotheses, our analysis suggested that both unilateral and bilateral stimulation impaired performance, considering that the performance in the stimulation conditions improved significantly less than the sham condition. Despite post-hoc indicating no significant difference between unilateral and bilateral, unilateral caused slightly lower impairment compared to bilateral.

## 3 Experiment 2 – dorsolateral prefrontal cortex stimulation

Another brain region which has been suggested to be involved in motor learning, and more specifically whole-body movement is the dorsolateral prefrontal cortex (DLPFC). The DLPFC is associated with working-memory (Techayusukcharoen, Iida and Aoki, 2019), long-term memory (Blumenfeld & Ranganath, 2006), attention (Kondo, Osaka & Osaka, 2004), planning (Kaller et al, 2011) and reasoning (Nelson et al. 2016). Subsequently, such aspects have also been heightened through the use of tDCS (Boggio et al. 2006; Fregni et al. 2005; Harty et al. 2014; Javadi & Walsh, 2012; Javadi, Cheng & Walsh, 2012). The DLPFC is important as it links visual cues with information within working memory to produce the required movement (Fuster, 2001). Additionally, Fujiyama (2016) suggests that the DLPFC is active in both preparation of movement but also control of movement. Specifically, the DLPFC is prominent when the task is both novel and the learning paradigm involves observation and imitation learning (Mineo et al. 2018). These aspects are important in the process of learning dance, therefore suggesting that the DLPFC may be an efficacious brain region for improving an individual’s ability in motor learning (Pascual-Leone, Wassermann, Grafman and Hallett, 1996). Additionally, it has been shown that tDCS over the DLPFC can modulate task switching and multitasking (Leite, Carvalho, Regni and Goncalves, 2011; Frank, Harty, Kluge & Kadosh, 2018; Hsu et al, 2015, Nelson et al. 2016), which are important for whole-body movement as individuals will need to think about both spatial awareness and coordination concordantly.

We hypothesised that active anodal tDCS over the left-DLPFC will be more beneficial than sham stimulation at improving an individual’s ability to learn dance moves. Furthermore, we hypothesised that bilateral anodal-tDCS will be more advantageous than unilateral stimulation at improving an individual’s ability to learn dance moves.

### 3.1 Method

#### 3.1.1 Participants

The sample included 31 participants (all females, age Mean[SD] = 19.32[1.33] years old). None of the participants had experience with Latin or Ballroom dancing, as indicated in a prestudy questionnaire.

#### 3.1.2 Materials

The same materials were used for both experiments.

*Transcranial Direct Current Stimulation (tDCS)* – One or two tDCS (NeuroConn, Germany) stimulators were used, depending on the protocol), with current amplitude of 1.5mA and 10 seconds ramp up and down. Stimulation was applied to either the left-DLPFC (Unilateral stimulation) or both the left- and right-DLPFC (Bilateral stimulation). According to the international EEG 10-20 system, the active anodal electrode was placed on F3 (Unilateral stimulation), and F3 and F4 (Bilateral stimulation) (Jasper, 1958). The cathode electrode was applied on the upper arm, contralateral to the side of the anode electrode. When the protocol required bilateral stimulation both upper arms were used, Figure 1. Electrodes were 5×5cm^2^, and were soaked in a salt-water solution. For sham stimulation, electrode placement was that of the unilateral condition, however stimulation was only applied for 10 seconds before being turned off.

#### 3.1.3 Procedure

The procedure used for the first experiment was repeated for the second experiment.

### 3.2 Results

An rANOVA was run with stimulation condition (Bilateral/Unilateral/Sham) and session (Post-intervention/Follow-up) as within subject factors on performance percentage change from baseline. Results showed a significant main effect of condition (*F*(2,46)= 7.517, *p* = 0.001, *η_p_^2^* = 0.246) and a significant main effect session (*F*(1,23)= 65.586, *p* < 0.001, *η_p_^2^* = 0.740), but a non-significant interaction effect (*F*(2,46)= 0.542, *p* = 0.585, *η_p_^2^* =0.023).

Post-hoc paired-sample t-tests showed a non-significant difference between Sham and Unilateral (Sham mean[SD] = 42.40[0.18]%, Unilateral 44.20[0.18]%, t(29) = 0.740, p = 0.298, d = 0.135), but a significant difference between Sham and Bilateral (Bilateral 34.20[0.14]%, t(29) = 2.562, p = 0.019, d = 0.468), and Unilateral and Bilateral (t(29) = 3.421, p = 0.003, d = 0.625). See Figure 4 for a summary of the performance of the participants.

**Figure 4.**
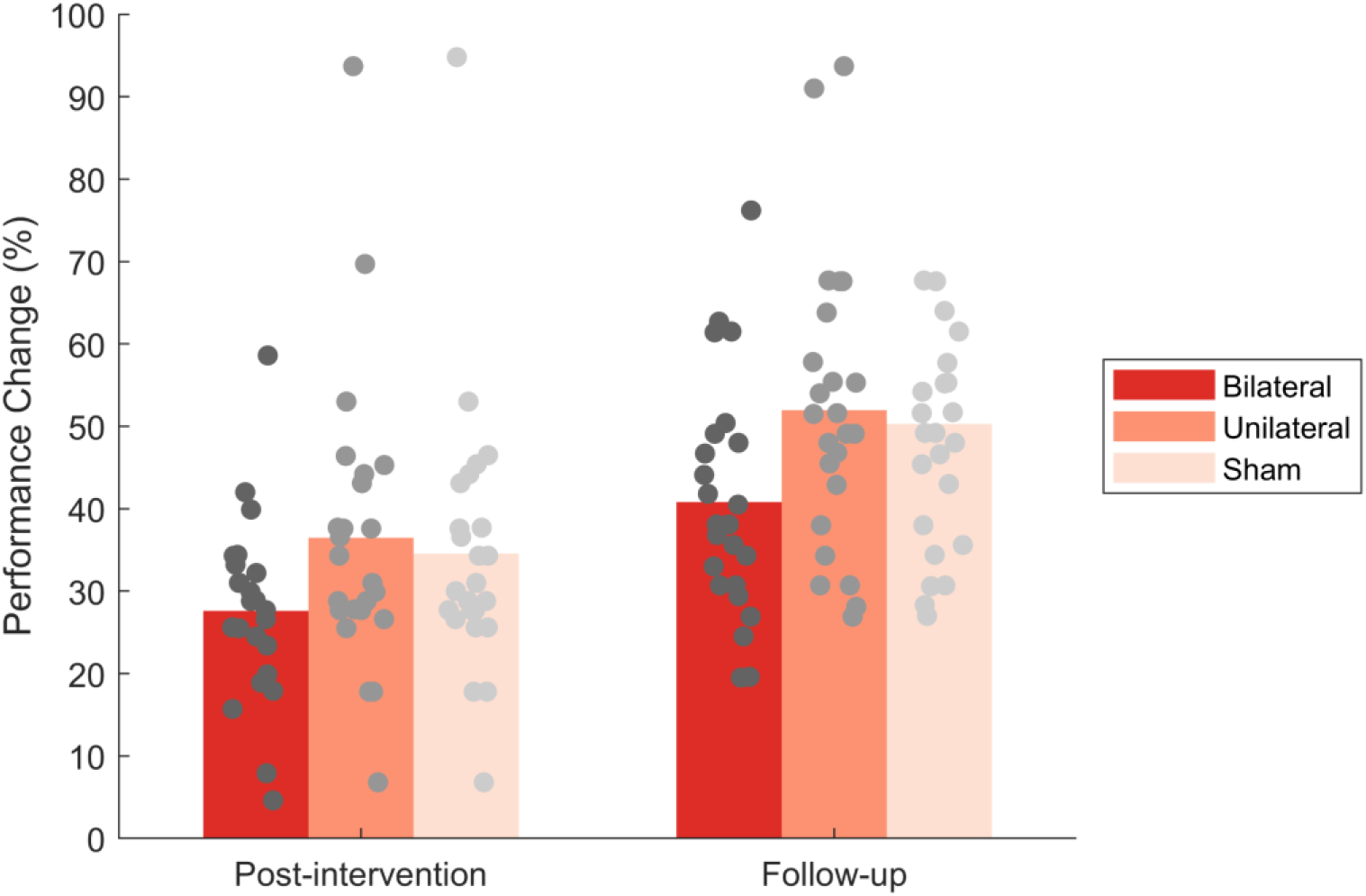
Performance percentage change from the baseline for the participants in different conditions and sessions in Experiment 2 with stimulation of the dorsolateral prefrontal cortex. Bilateral stimulation showed significantly lower performance compared to Sham *(p* = 0.019) and Unilateral *(p* = 0.003) stimulation conditions. No significant difference was found between Sham and Unilateral stimulation conditions *(p* = 0.298).

### 3.3 Summary

In the second experiment, we investigated the effects of unilateral anodal-tDCS over the left-DLPFC and bilateral anodal-tDCS over both DLPFC, on the learning of Ballroom and Latin dance moves. Similar to the first experiment, we expected that active stimulation would be more effective than sham stimulation in improving dance performance and that bilateral would be more effective than unilateral. Contrary to our hypotheses, however, it was found that unilateral tDCS did not significantly improve performance compared to the Sham condition. Furthermore, bilateral stimulation impaired performance as compared to the other conditions.

## 4 Discussion

In this study, we investigated the effect of tDCS on motor behaviour that requires whole-body movement. Stimulation was delivered either unilaterally or bilaterally over either the primary motor cortex (PMC) or dorsolateral prefrontal cortex (DLPFC). Our results showed that bilateral stimulation impaired performance as compared to the sham stimulation, regardless of area of stimulation. Unilateral stimulation showed impairing effects only if applied to the right-PMC. Unilateral stimulation of the left-DLPFC did not differ significantly from the sham stimulation.

There is an abundance of research, which has successfully found a positive effect of tDCS on motor movement and learning (Kang, Summers & Cauraugh, 2016; Nitsche et al. 2003; Reis & Fritsch, 2011). These studies, however, demonstrated learning effects through simple motor tasks, which have very limited ecological validity (Ronsse et al. 2009). Every day activities are dependent on complex whole-body motor movement, multi-tasking and an awareness of direction in space and speed (Bläsing, Calvo-Merino, Cross & Jola, 2012). Our results showed impairing effects of bilateral tDCS over PMC and DLPFC, and also impairing effect of unilateral tDCS over PMC. Therefore, special considerations need to be made in order to harness beneficial effects of tDCS. A few mechanisms could explain the impairing effects observed in this study.

Effects of tDCS on motor performance is contingent on the complexity of the task. Previous research has found that PMC stimulation does not improve performance on bimanual or complex tasks (Fleming et al, 2017; Furuya, Klaus, Nitsche, Paulus & Altenmüller, 2014; Mesquita, Lage, Franchini, Romano-Silva & Alberquerque, 2019; Pixa et al, 2019), which are cognitively more demanding (Szameitat, Lesien, von Cramon, Sterr & Schubert, 2006). Complex motor tasks engage brain networks beyond networks engaged in simple motor tasks. Therefore, stimulation protocols used for simple motor movements might not be suitable for complex whole-body movements (Pixa & Pollok, 2018). For complex whole-body movements, individuals not only need to coordinate their body parts, but they also need to attend to and organise multiple pieces of information (Brown, Martinez & Parsons 2005). Additionally, research has suggested that the modulation of one region is not appropriate for complex wholebody movement (Fischer et al. 2017; Pixa & Pollok, 2018; Vancleef et al, 2016). Therefore, while stimulation of the motor cortex might help with motor movement, and stimulation of the DLPFC might help with information processing, isolated stimulation of these brain areas might not be able to drive complex brain networks required in complex whole-body movements such as dance.

Contrary to the majority of past research, our results showed that bilateral tDCS in both experiments led to impaired performance. One possible explanation is that the effects of tDCS might reverse depending on the task. Bortoletto, Pellicciari, Rodella and Miniussi (2014) found that when anodal-tDCS of the right-PMC was paired with a fast motor learning protocol compared to a slow motor learning protocol, learning and performance was reduced. Authors suggested that the learning of a fast motor task increases cortical excitability alone and that in addition to the excitatory effects of tDCS lead to reversal of the facilitatory effects. According to the neuronal-noise framework the effects of tDCS are dependent on the strength of the signal in relation to the amount of noise present (Miniussi, Harris & Ruzzoli, 2013). Signal relates to neural activity operational to the task and the noise conveys random neural activity.

tDCS effects are state-dependant (Hsu, Juan & Tseng, 2016). Subsequently, Pixa and Pollok, (2017) suggest that during complex movement there is increased activity in prefrontal, parietal and temporal areas. Hence because neurones are highly active due to motor practice the noise levels will increase, subsequently decreasing the signal-to-noise ratio and impairing performance. Additionally, Miniussi, Harris and Ruzzoli (2013) found that the reversibility effects of tDCS are also seen when tasks, which require skill but are not established, are combined with tDCS. As motor-tasks become more established neuronal noise levels decrease, leaving the signal to clearly materialise and allow anodal-tDCS to enable performance improvement. For participants within our study, Ballroom and Latin dance was a novel task. So, it can be expected that neuronal noise levels were high. Therefore, any tDCS applied would have impaired learning and performance of dance. Therefore, it could be suggested that by practicing more and allowing the task to become more habitual the signal will be able to materialise more clearly allowing tDCS to improve behaviour.

Another possible explanation for the impairing effects of bilateral tDCS in our study is that we stimulated both lateralities anodally, while the majority of past research applying bilateral tDCS used an incongruent montage; for example, anodal-tDCS of the right-PMC and cathodal-tDCS of the left-PMC. This incongruent montage has proved beneficial in improving; dual task performance (Ljubisavljevic et al, 2019), motor learning (Karok & Witney, 2013), fatigue in fast motor tasks (Arias et al, 2016) and rehabilitation for stroke patients (Goodwill et al, 2016; Lefebvre et al, 2013). It has, however, shown that incongruent bilateral tDCS favours one laterality over the other. Javadi et al (2015) showed that anodal- and cathodal-tDCS of the right- and left-PMC, respectively, leads to increased and decreased response on the left- and right-hand side of the body, respectively. Inversely, the opposite polarity of stimulation led to the opposite effect. Similarly, research using incongruent montages, use this montage to solely modulate a specific limb (Arias et al, 2016; Karok, Fletcher & Witney, 2017; Mordillo-Mateos et al, 2012; Naros et al, 2016; Vines et al, 2008). The success of this tDCS protocol is suggested to be due to the reduction of interhemispheric inhibition. This idea of decreasing the excitability of one hemisphere to promote the excitability of the other hemisphere is logical when we are improving unimanual skills, where cortical excitability is localised to one area (Gomes-Osman & Field-Fote, 2013). But with whole-body and complex movements balanced bilateral activation and effective communication between the hemispheres is required (Chettouf et al, 2020; Waller et al, 2008). Therefore, we opted not to use incongruent stimulation and instead apply anodal-tDCS bilaterally.

Previous research has found success with anodal bilateral stimulation. Angius et al (2018) showed that with anodal bilateral tDCS, participants had increased endurance in a cycling task. Hadoush et al (2018) suggested that bilateral anodal-tDCS to the DLPFC and PMC, concordantly, improved balance and functional ability and in turn reduced fear of falling. And Gomes-Osman and Field-Fote (2013), found that after bilateral anodal-tDCS, participants performed better on a bimanual typing task. However, the nature of these tasks is simple requiring minimal cognitive input to perform the tasks successfully. Therefore, the comparability to this study is reduced. Subsequently, Mequita et al (2019), also showed a deleterious effect of anodal bilateral tDCS, within a taekwondo task. This task is similar to the dance task, involved in this study, in that it required high complexity involving multi-joint actions and awareness of self in space. Consequently, this reaffirms the importance of considering task complexity when modulating motor behaviour with tDCS.

A possible limitation of this study is that we did not measure the adverse effects of the stimulation on participants. Participants were blinded to type of stimulation they were receiving, however this was not tested. It is known that tDCS can cause mild irritation. Subsequently with bilateral tDCS there would have been more irritation than with unilateral stimulation. Therefore, the increased irritation, with bilateral tDCS, may have made it harder for participants to focus on learning the dance moves. Accordingly, it would be useful for future research to evaluate the effect of tDCS irritation has on learning motor tasks.

Additionally, stimulation would not have been focal (see Figure 1). The size of the electrodes (25cm^2^) would result in other areas of the brain being stimulated, regions which may not be crucial in the enhancement of motor learning, or may be inhibitory to motor learning. Bastani and Jaberzadeh (2013) found that 12cm^2^ electrodes increased corticospinal activity the most compared to 25cm^2^ and 35cm^2^ electrodes. Thus, using smaller electrodes could increase the efficacy of stimulating the primary region of choice.

When using tDCS for clinical purposes, a unilateral or incongruent montage is most often used (Kim et al, 2019; Lefebvre et al, 2013). Primarily in clinical studies which use tDCS, the purpose is to improve performance only on the affected side of the brain (Bolognini et al, 2011; Raithatha et al, 2016; Yozbatiran et al, 2016). Both unilateral and incongruent bilateral tDCS can offer explanations to how this process happens. Contrarily, as research using a congruent bilateral montage is much rarer there are many unknowns about the mechanistic effects. The same can additionally be said for complex motor movement, where the neural underpinnings are still uncertain. Therefore, to apply congruent tDCS to a clinical population without knowing how it will affect the neural mechanisms is unwise as a further detriment to their movement could be made.

The initial literature suggested that performance effects are heavily influenced by an individual’s baseline performance. Whilst this study made sure that all participants did not have Ballroom and Latin dance experience, there would still have been differences in baseline score due to individual’s differing in speed of learning, coordination and rhythm. Subsequently, it would be able to see if participants who had lower initial baseline scores benefited from the tES protocol more than individuals with higher baseline scores.

To summarise, when we are looking at modulating brain regions in association with motor learning, complexity of the task needs to be considered. Furthermore, research needs to look more closely about the neural mechanisms that underpin complex whole-body movement and how such mechanisms can be modulated. While there is a great body of literature on the improving effects of tDCS on simple motor learning tasks, tDCS in combination with complex whole-body movements should be considered cautiously.

## Supplementary Material

Dance stimuli and scoring sheets can be found on http://doi.org/10.17605/OSF.IO/VSA36.

## Acknowledgements

Authors would like to thank Michael Willcox for his help in preparing the stimuli, and Laura Smith, John Allen, Adam Britcher and Frank Gasking for their help in setting up the experiment and technical support.

## Notes

http://doi.org/10.17605/OSF.IO/VSA36

### Competing Interest Statement

The authors have declared no competing interest.

## References

Allman, C., Amadi, U., Winkler, A. M., Wilkins, L., Filippini, N., Kischka, U., … & Johansen-Berg, H. (2016). Ipsilesional anodal tDCS enhances the functional benefits of rehabilitation in patients after stroke. Science translational medicine, 8(330), 330re1–330re1.

Angius, L., Mauger, A. R., Hopker, J., Pascual-Leone, A., Santarnecchi, E., & Marcora, S. M. (2018). Bilateral extracephalic transcranial direct current stimulation improves endurance performance in healthy individuals. Brain stimulation, 11(1), 108–117.

Apšvalka, D., Ramsey, R., & Cross, E. S. (2018). Anodal tDCS over primary motor cortex provides no advantage to learning motor sequences via observation. Neural plasticity, 2018.

Arias, P., Corral-Bergantiños, Y., Robles-García, V., Madrid, A., Oliviero, A., & Cudeiro, J. (2016). Bilateral tDCS on primary motor cortex: effects on fast arm reaching tasks. PLoS One, 11(8), e0160063.

Arnao, V., Riolo, M., Carduccio, F., Tuttolomondo, A., D’Amelio, M., Brighina, F., … & Aridon, P. (2019). Effects of transcranial random noise stimulation combined with Graded Repetitive Arm Supplementary Program (GRASP) on motor rehabilitation of the upper limb in sub-acute ischemic stroke patients: a randomized pilot study. Journal of Neural Transmission, 126(12), 1701–1706.

Bindman, L. J., Lippold, O. C. J., & Redfearn, J. W. T. (1964). The action of brief polarizing currents on the cerebral cortex of the rat (1) during current flow and (2) in the production of long-lasting after-effects. The Journal of physiology, 172(3), 369.

Bläsing, B., Calvo-Merino, B., Cross, E. S., Jola, C., Honisch, J., & Stevens, C. J. (2012). Neurocognitive control in dance perception and performance. Acta psychologica, 139(2}, 300–308.

Blumenfeld, R. S., & Ranganath, C. (2006). Dorsolateral prefrontal cortex promotes longterm memory formation through its role in working memory organization. Journal of Neuroscience, 26(3), 916–925.

Boggio, P. S., Ferrucci, R., Rigonatti, S. P., Covre, P., Nitsche, M., Pascual-Leone, A., & Fregni, F. (2006). Effects of transcranial direct current stimulation on working memory in patients with Parkinson’s disease. Journal of the neurological sciences, 249(1), 31–38.

Bolognini, N., Vallar, G., Casati, C., Latif, L. A., El-Nazer, R., Williams, J., … & Fregni, F. (2011). Neurophysiological and behavioral effects of tDCS combined with constraint-induced movement therapy in poststroke patients. Neurorehabilitation and neural repair, 25(9), 819–829.

Bortoletto, M., Pellicciari, M. C., Rodella, C., & Miniussi, C. (2015). The interaction with task-induced activity is more important than polarization: a tDCS study. Brain stimulation, 8(2), 269–276.

Brown, S., Martinez, M. J., & Parsons, L. M. (2006). The neural basis of human dance. Cerebral cortex, 16(8), 1157–1167.

Ciechanski, P., & Kirton, A. (2016). Transcranial direct-current stimulation (tDCS): principles and emerging applications in children. In Pediatric Brain Stimulation (pp. 85–115). Academic Press.

Chettouf, S., Rueda-Delgado, L. M., de Vries, R., Ritter, P., & Daffertshofer, A. (2020). Are unimanual movements bilateral?. Neuroscience & Biobehavioral Reviews.

Cuypers, K., Leenus, D. J., van den Berg, F. E., Nitsche, M. A., Thijs, H., Wenderoth, N., & Meesen, R. L. (2013). Is motor learning mediated by tDCS intensity?. PLoS One, 8(6), e67344.

Dumel, G., Bourassa, M. È., Charlebois-Plante, C., Desjardins, M., Doyon, J., Saint-Amour, D., & De Beaumont, L. (2018). Multisession anodal transcranial direct current stimulation induces motor cortex plasticity enhancement and motor learning generalization in an aging population. Clinical Neurophysiology, 129(2), 494–502.

Duncan, R. P., & Earhart, G. M. (2014). Are the effects of community-based dance on Parkinson disease severity, balance, and functional mobility reduced with time? A 2-year prospective pilot study. The Journal of Alternative and Complementary Medicine, 20(10), 757–763.

Dushanova, J., & Donoghue, J. (2010). Neurons in primary motor cortex engaged during action observation. European Journal of Neuroscience, 31(2), 386–398.

Ehsani, F., Bakhtiary, A. H., Jaberzadeh, S., Talimkhani, A., & Hajihasani, A. (2016). Differential effects of primary motor cortex and cerebellar transcranial direct current stimulation on motor learning in healthy individuals: a randomized double-blind sham-controlled study. Neuroscience research, 112, 10–19.

Filar-Mierzwa, K., Długosz, M., Marchewka, A., Dąbrowski, Z., & Poznańska, A. (2017). The effect of dance therapy on the balance of women over 60 years of age: The influence of dance therapy for the elderly. Journal of women & aging, 29(4), 348–355.

Fischer, D. B., Fried, P. J., Ruffini, G., Ripolles, O., Salvador, R., Banus, J., … & Fox, M. D. (2017). Multifocal tDCS targeting the resting state motor network increases cortical excitability beyond traditional tDCS targeting unilateral motor cortex. Neuroimage, 157, 34–44.

Fleming, M. K., Rothwell, J. C., Sztriha, L., Teo, J. T., & Newham, D. J. (2017). The effect of transcranial direct current stimulation on motor sequence learning and upper limb function after stroke. Clinical Neurophysiology, 128(7), 1389–1398.

Frank, B., Harty, S., Kluge, A., & Cohen Kadosh, R. (2018). Learning while multitasking: short and long-term benefits of brain stimulation. Ergonomics, 61(11), 1454–1463.

Fregni, F., Boggio, P. S., Nitsche, M., Bermpohl, F., Antal, A., Feredoes, E., … & Pascual-Leone, A. (2005). Anodal transcranial direct current stimulation of prefrontal cortex enhances working memory. Experimental brain research, 166(1), 23–30.

Fujiyama, H., Van Soom, J., Rens, G., Cuypers, K., Heise, K. F., Levin, O., & Swinnen, S. P. (2016). Performing two different actions simultaneously: The critical role of interhemispheric interactions during the preparation of bimanual movement. Cortex, 77, 141–154.

Furuya, S., Klaus, M., Nitsche, M. A., Paulus, W., & Altenmüller, E. (2014). Ceiling effects prevent further improvement of transcranial stimulation in skilled musicians. Journal of Neuroscience, 34(41), 13834–13839.

Fuster, J. M. (2001). The prefrontal cortex—an update: time is of the essence. Neuron, 30(2), 319–333.

Giustiniani, A., Tarantino, V., Bonaventura, R. E., Smirni, D., Turriziani, P., & Oliveri, M. (2019). Effects of low-gamma tACS on primary motor cortex in implicit motor learning. Behavioural brain research, 376, 112170.

Gomes-Osman, J., & Field-Fote, E. C. (2013). Bihemispheric anodal corticomotor stimulation using transcranial direct current stimulation improves bimanual typing task performance. Journal of motor behavior, 45(4), 361–367.

Goodwill, A. M., Teo, W. P., Morgan, P., Daly, R. M., & Kidgell, D. J. (2016). Bihemispheric-tDCS and upper limb rehabilitation improves retention of motor function in chronic stroke: a pilot study. Frontiers in human neuroscience, 10, 258.

Hadoush, H., Al-Jarrah, M., Khalil, H., Al-Sharman, A., & Al-Ghazawi, S. (2018). Bilateral anodal transcranial direct current stimulation effect on balance and fearing of fall in patient with Parkinson’s disease. NeuroRehabilitation, 42(1), 63–68.

Harty, S., Robertson, I. H., Miniussi, C., Sheehy, O. C., Devine, C. A., McCreery, S., & O’Connell, R. G. (2014). Transcranial direct current stimulation over right dorsolateral prefrontal cortex enhances error awareness in older age. Journal of Neuroscience, 34(10), 3646–3652.

Hashemirad, F., Zoghi, M., Fitzgerald, P. B., & Jaberzadeh, S. (2016). The effect of anodal transcranial direct current stimulation on motor sequence learning in healthy individuals: a systematic review and meta-analysis. Brain and cognition, 102, 1–12.

Honda, M., Wise, S. P., Weeks, R. A., Deiber, M. P., & Hallett, M. (1998). Cortical areas with enhanced activation during object-centred spatial information processing. A PET study. Brain: a journal of neurology, 121(11), 2145–2158.

Hoy, K. E., Emonson, M. R., Arnold, S. L., Thomson, R. H., Daskalakis, Z. J., & Fitzgerald, P. B. (2013). Testing the limits: investigating the effect of tDCS dose on working memory enhancement in healthy controls. Neuropsychologia, 51(9), 1777–1784.

Hsu, W. Y., Zanto, T. P., Anguera, J. A., Lin, Y. Y., & Gazzaley, A. (2015). Delayed enhancement of multitasking performance: effects of anodal transcranial direct current stimulation on the prefrontal cortex. Cortex, 69, 175–185.

Hsu, T. Y., Juan, C. H., & Tseng, P. (2016). Individual differences and state-dependent responses in transcranial direct current stimulation. Frontiers in human neuroscience, 10, 643.

Huang, Y., Datta, A., Bikson, M., & Parra, L. C. (2019). Realistic volumetric-approach to simulate transcranial electric stimulation—ROAST—a fully automated open-source pipeline. Journal of Neural Engineering, 16(5), 056006. https://doi.org/10.1088/1741-2552/ab208d

Jackson, A. K., de Albuquerque, L. L., Pantovic, M., Fischer, K. M., Guadagnoli, M. A., Riley, Z. A., & Poston, B. (2019). Cerebellar transcranial direct current stimulation enhances motor learning in a complex overhand throwing task. The Cerebellum, 18(4), 813–816.

Jasper, H. (1958). Report of the committee on methods of clinical examination in electroencephalography. Electroencephalogr Clin Neurophysiol, 10, 370–375.

Javadi, A.-H., & Walsh, V. (2012). Transcranial direct current stimulation (tDCS) of the left dorsolateral prefrontal cortex modulates declarative memory. Brain stimulation, 5(3), 231–241.

Javadi, A.-H., Cheng, P., & Walsh, V. (2012). Short duration transcranial direct current stimulation (tDCS) modulates verbal memory. Brain stimulation, 5(4), 468–474.

Javadi, A.-H., Beyko, A., Walsh, V., & Kanai, R. (2015). Transcranial direct current stimulation of the motor cortex biases action choice in a perceptual decision task. Journal of cognitive neuroscience, 27(11), 2174–2185.

Kakei, S., Hoffman, D. S., & Strick, P. L. (1999). Muscle and movement representations in the primary motor cortex. Science, 285(5436), 2136–2139.

Kaller, C. P., Heinze, K., Frenkel, A., Läppchen, C. H., Unterrainer, J. M., Weiller, C., … & Rahm, B. (2013). Differential impact of continuous theta-burst stimulation over left and right DLPFC on planning. Human brain mapping, 34(1), 36–51.

Kami, A. T., Sadler, C., Nantel, J., & Carlsen, A. N. (2018). Transcranial direct current stimulation (TDCS) over supplementary motor area (SMA) improves upper limb movement in individuals with Parkinson’s disease. Journal of Exercise, Movement, and Sport (SCAPPS refereed abstracts repository), 50(1), 36–36.

Kaminski, E., Hoff, M., Sehm, B., Taubert, M., Conde, V., Steele, C. J., … & Ragert, P. (2013). Effect of transcranial direct current stimulation (tDCS) during complex whole body motor skill learning. Neuroscience letters, 552, 76–80.

Kaminski, E., Steele, C. J., Hoff, M., Gundlach, C., Rjosk, V., Sehm, B., … & Ragert, P. (2016). Transcranial direct current stimulation (tDCS) over primary motor cortex leg area promotes dynamic balance task performance. Clinical Neurophysiology, 127(6), 2455–2462.

Kang, N., Summers, J. J., & Cauraugh, J. H. (2016). Transcranial direct current stimulation facilitates motor learning post-stroke: a systematic review and meta-analysis. Journal of Neurology, Neurosurgery & Psychiatry, 87(4), 345–355.

Karok, S., & Witney, A. G. (2013). Enhanced motor learning following task-concurrent dual transcranial direct current stimulation. PloS one, 8(12), e85693.

Karok, S., Fletcher, D., & Witney, A. G. (2017). Task-specificity of unilateral anodal and dual-M1 tDCS effects on motor learning. Neuropsychologia, 94, 84–95.

Kim, W. S., Lee, K., Kim, S., Cho, S., & Paik, N. J. (2019). Transcranial direct current stimulation for the treatment of motor impairment following traumatic brain injury. Journal of NeuroEngineering and Rehabilitation, 16(1), 1–10.

Kida, H., & Mitsushima, D. (2018). Mechanisms of motor learning mediated by synaptic plasticity in rat primary motor cortex. Neuroscience Research, 128, 14–18.

Kondo, H., Osaka, N., & Osaka, M. (2004). Cooperation of the anterior cingulate cortex and dorsolateral prefrontal cortex for attention shifting. Neuroimage, 23(2), 670–679.

Lakes, K. D., Marvin, S., Rowley, J., San Nicolas, M., Arastoo, S., Viray, L., … & Jurnak, F. (2016). Dancer perceptions of the cognitive, social, emotional, and physical benefits of modern styles of partnered dancing. Complementary therapies in medicine, 26, 117–122.

Lefebvre, S., Laloux, P., Peeters, A., Desfontaines, P., Jamart, J., & Vandermeeren, Y. (2013). Dual-tDCS enhances online motor skill learning and long-term retention in chronic stroke patients. Frontiers in human neuroscience, 6, 343.

Leite, J., Carvalho, S., Fregni, F., & Gonçalves, O. F. (2011). Task-specific effects of tDCS-induced cortical excitability changes on cognitive and motor sequence set shifting performance. PloS one, 6(9), e24140.

Lindenberg, R., Renga, V., Zhu, L. L., Nair, D., & Schlaug, G. M. D. P. (2010). Bihemispheric brain stimulation facilitates motor recovery in chronic stroke patients. Neurology, 75(24), 2176–2184.

Lindenberg, R., Sieg, M. M., Meinzer, M., Nachtigall, L., & Flöel, A. (2016). Neural correlates of unihemispheric and bihemispheric motor cortex stimulation in healthy young adults. Neuroimage, 140, 141–149.

Ljubisavljevic, M. R., Joji, O., Filipovic, S., Bjekic, J., Szolics, M., & Nagelkerke, N. (2019). Effects of tDCS of Dorsolateral Prefrontal Cortex on Dual-Task Performance Involving Manual Dexterity and Cognitive Task in Healthy Older Adults. Frontiers in aging neuroscience, 11, 144.

McCambridge, A. B., Bradnam, L. V., Stinear, C. M., & Byblow, W. D. (2011). Cathodal transcranial direct current stimulation of the primary motor cortex improves selective muscle activation in the ipsilateral arm. Journal of Neurophysiology, 105(6), 2937–2942.

Meng, X., Li, G., Jia, Y., Liu, Y., Shang, B., Liu, P., … & Chen, L. (2020). Effects of dance intervention on global cognition, executive function and memory of older adults: a meta-analysis and systematic review. Aging clinical and experimental research, 32(1), 7–19.

Mesquita, P. H. C., Lage, G. M., Franchini, E., Romano-Silva, M. A., & Albuquerque, M. R. (2019). Bi-hemispheric anodal transcranial direct current stimulation worsens taekwondo-related performance. Human movement science, 66, 578–586.

Mineo, L., Fetterman, A., Concerto, C., Warren, M., Infortuna, C., Freedberg, D., … & Battaglia, F. (2018). Motor facilitation during observation of implied motion: Evidence for a role of the left dorsolateral prefrontal cortex. International Journal of Psychophysiology, 128, 47–51.

Miniussi, C., Harris, J. A., & Ruzzoli, M. (2013). Modelling non-invasive brain stimulation in cognitive neuroscience. Neuroscience & Biobehavioral Reviews, 37(8), 1702–1712.

Mordillo-Mateos, L., Turpin-Fenoll, L., Millán-Pascual, J., Núñez-Pérez, N., Panyavin, I., Gómez-Argüelles, J. M., … & Oliviero, A. (2012). Effects of simultaneous bilateral tDCS of the human motor cortex. Brain stimulation, 5(3), 214–222.

Moura, R. C. F., Santos, C., Collange Grecco, L., Albertini, G., Cimolin, V., Galli, M., & Oliveira, C. (2017). Effects of a single session of transcranial direct current stimulation on upper limb movements in children with cerebral palsy: A randomized, sham-controlled study. Developmental neurorehabilitation, 20(6), 368–375.

Muellbacher, W., Ziemann, U., Boroojerdi, B., Cohen, L., & Hallett, M. (2001). Role of the human motor cortex in rapid motor learning. Experimental brain research, 136(4), 431–438.

Naros, G., Geyer, M., Koch, S., Mayr, L., Ellinger, T., Grimm, F., & Gharabaghi, A. (2016). Enhanced motor learning with bilateral transcranial direct current stimulation: impact of polarity or current flow direction?. Clinical Neurophysiology, 127(4), 2119–2126.

Nelson, J., McKinley, R. A., Phillips, C., McIntire, L., Goodyear, C., Kreiner, A., & Monforton, L. (2016). The effects of transcranial direct current stimulation (tDCS) on multitasking throughput capacity. Frontiers in Human Neuroscience, 10, 589.

Nitsche, M. A., & Paulus, W. (2001). Sustained excitability elevations induced by transcranial DC motor cortex stimulation in humans. Neurology, 57(10), 1899–1901.

Nitsche, M. A., Schauenburg, A., Lang, N., Liebetanz, D., Exner, C., Paulus, W., & Tergau, F. (2003). Facilitation of implicit motor learning by weak transcranial direct current stimulation of the primary motor cortex in the human. Journal of cognitive neuroscience, 15(4), 619–626.

Parkin, B. L., Ekhtiari, H., & Walsh, V. (2015). Non-invasive Human Brain Stimulation in Cognitive Neuroscience: A Primer. Neuron, 87(5), 932–945. https://doi.org/10.1016/j.neuron.2015.07.032

Parkin, B. L., Bhandari, M., Glen, J. C., & Walsh, V. (2019). The physiological effects of transcranial electrical stimulation do not apply to parameters commonly used in studies of cognitive neuromodulation. Neuropsychologia, 128, 332–339.

Pascual-Leone, A., Wassermann, E. M., Grafman, J., & Hallett, M. (1996). The role of the dorsolateral prefrontal cortex in implicit procedural learning. Experimental brain research, 107(3), 479–485.

Pixa, N. H., & Pollok, B. (2018). Effects of tDCS on bimanual motor skills: a brief review. Frontiers in behavioral neuroscience, 12, 63.

Pixa, N. H., Berger, A., Steinberg, F., & Doppelmayr, M. (2019). Parietal, but not motor cortex, HD-atDCS deteriorates learning transfer of a complex bimanual coordination task. Journal of Cognitive Enhancement, 3(1), 111–123.

Raithatha, R., Carrico, C., Powell, E. S., Westgate, P. M., Chelette II, K. C., Lee, K., … & Sawaki, L. (2016). Non-invasive brain stimulation and robot-assisted gait training after incomplete spinal cord injury: A randomized pilot study. NeuroRehabilitation, 38(1), 15–25.

Reis, J., & Fritsch, B. (2011). Modulation of motor performance and motor learning by transcranial direct current stimulation. Current opinion in neurology, 24(6), 590–596.

Rocha, S., Silva, E., Foerster, Á., Wiesiolek, C., Chagas, A. P., Machado, G., … & Monte-Silva, K. (2016). The impact of transcranial direct current stimulation (tDCS) combined with modified constraint-induced movement therapy (mCIMT) on upper limb function in chronic stroke: a double-blind randomized controlled trial. Disability and rehabilitation, 38(7), 653–660.

Ronsse, R., Miall, R. C., & Swinnen, S. P. (2009). Multisensory integration in dynamical behaviors: maximum likelihood estimation across bimanual skill learning. Journal of Neuroscience, 29(26) 8419–8428.

Sehm, B., Kipping, J. A., Schäfer, A., Villringer, A., & Ragert, P. (2013). A comparison between uni-and bilateral tDCS effects on functional connectivity of the human motor cortex. Frontiers in human neuroscience, 7, 183.

Stagg, C. J., Jayaram, G., Pastor, D., Kincses, Z. T., Matthews, P. M., & Johansen-Berg, H. (2011). Polarity and timing-dependent effects of transcranial direct current stimulation in explicit motor learning. Neuropsychologia, 49(5), 800–804.

Steiner, K. M., Enders, A., Thier, W., Batsikadze, G., Ludolph, N., Ilg, W., & Timmann, D. (2016). Cerebellar tDCS does not improve learning in a complex whole body dynamic balance task in young healthy subjects. PloS one, 11(9), e0163598.

Szameitat, A. J., Lepsien, J., Von Cramon, D. Y., Sterr, A., & Schubert, T. (2006). Taskorder coordination in dual-task performance and the lateral prefrontal cortex: an event-related fMRI study. Psychological research, 70(6), 541–552.

Talimkhani, A., Abdollahi, I., Mohseni-Bandpei, M. A., Ehsani, F., Khalili, S., & Jaberzadeh, S. (2019). Differential effects of unihemispheric concurrent dual-site and conventional tDCS on motor learning: A randomized, sham-controlled study. Basic and clinical neuroscience, 10(1), 59.

Techayusukcharoen, R., Iida, S., & Aoki, C. (2019). Observing brain function via functional near-infrared spectroscopy during cognitive program training (dual task) in young people. Journal of physical therapy science, 31(7), 550–555.

Vancleef, K., Meesen, R., Swinnen, S. P., & Fujiyama, H. (2016). tDCS over left M1 or DLPFC does not improve learning of a bimanual coordination task. Scientific reports, 6, 35739.

Vines, B. W., Cerruti, C., & Schlaug, G. (2008). Dual-hemisphere tDCS facilitates greater improvements for healthy subjects’ non-dominant hand compared to uni-hemisphere stimulation. BMC neuroscience, 9(1), 103.

Waller, S. M., Forrester, L., Villagra, F., & Whitall, J. (2008). Intracortical inhibition and facilitation with unilateral dominant, unilateral nondominant and bilateral movement tasks in left-and right-handed adults. Journal of the neurological sciences, 269(1-2), 96–104.

Waters-Metenier, S., Husain, M., Wiestler, T., & Diedrichsen, J. (2014). Bihemispheric transcranial direct current stimulation enhances effector-independent representations of motor synergy and sequence learning. Journal of Neuroscience, 34(3), 1037–1050.

Weightman, M., Brittain, J. S., Punt, D., Miall, R. C., & Jenkinson, N. (2020). Targeted tDCS selectively improves motor adaptation with the proximal and distal upper limb. Brain Stimulation.

Yamaguchi, T., Moriya, K., Tanabe, S., Kondo, K., Otaka, Y., & Tanaka, S. (2020). Transcranial direct-current stimulation combined with attention increases cortical excitability and improves motor learning in healthy volunteers. Journal of neuroengineering and rehabilitation, 17(1), 1–13.

Yozbatiran, N., Keser, Z., Davis, M., Stampas, A., O’Malley, M., Cooper-Hay, C., … & Francisco, G. E. (2016). Transcranial direct current stimulation (tDCS) of the primary motor cortex and robot-assisted arm training in chronic incomplete cervical spinal cord injury: a proof of concept sham-randomized clinical study. NeuroRehabilitation, 39(3), 401–411.

Zandvliet, S. B., Meskers, C. G., Kwakkel, G., & van Wegen, E. E. (2018). Short-term effects of cerebellar tDCS on standing balance performance in patients with chronic stroke and healthy age-matched elderly. The Cerebellum, 17(5), 575–589.

